# Equip *Fusobacterium nucleatum* genetic tool kits with compatible shuttle vectors and engineered intermediatory *E. coli* strains for enhanced transformation efficiency

**DOI:** 10.1101/2024.07.17.603877

**Authors:** Ling Liu, Yuzhang He, Tingting Zhang, Rui Geng, Yongmei Hu, Mingyue Luo, Hongwei Zhou, Xue Liu

## Abstract

*Fusobacterium nucleatum*, an oral microbe, is implicated in various human diseases, including oral-related diseases and tumors. However, efficient transformation was only achieved in limited strains of this bacterium. The challenges in conducting molecular level investigations of most strains due to their genetic intractability have hindered the biological studies of *F. nucleatum*. The restriction-modification (RM) systems is one of the known obstacles for efficient DNA transformation. Here, we used single molecule real time sequencing to elucidate the RM recognition sites and the corresponding methyltransferases (MTases) in two *F. nucleatum* strains. Based on the identified MTases, we engineered intermediatory *E. coli* host strains to bypass the RM systems, and showed that the plasmids harbored by these intermediatory strains can be efficiently electro-transformed, reaching 5000 transformants per microgram plasmids, paving the way for the development of efficient genetic modification tools. Furthermore, we successfully demonstrated that the conjugation-based DNA delivery to *F. nucleatum* can bypass the requirement of MTase methylations. By exploring the native plasmids from *F. nucleatum*, we identified new backbones for construction of shuttle vectors and established a dual-plasmid system for the first time, offering new avenues for genetic manipulation in this bacterium. Additionally, we evaluate promoters with variable strengths with a luciferase-based reporter system in *F. nucleatum*, providing valuable insights for future gene editing studies in bacterium and contributing to our understanding of its pathogenesis. All the tools developed in this study was shared via the WeKwikgene (https://wekwikgene.wllsb.edu.cn/).

**Impact Statement:** *Fusobacterium nucleatum*, a key opportunistic pathogen implicated in periodontal diseases, rheumatoid arthritis, and tumors, presents significant challenges due to its limited transformation efficiency and lack of gene-editing tools. In this study, we present an advancement -a streamlined and robust pipeline that enhances transformation efficiency by approximately 10^3^-fold in *F. nucleatum*, reaching 5000 CFU per microgram plasmids. This represents a significant breakthrough, marking the first report to achieve such a remarkable improvement in transformation efficiency in this pathogen. This improvement paves the way for the genome-wide level mutagenesis study in this bacterium.

## Introduction

The oral microbe *Fusobacterium nucleatum* is a gram-negative, anaerobic bacterium which gained increasing attention due to its association with human tumor, particularly the colorectal cancer (1–5). Our and others study provide rich evidence supporting the correlations of *Fusobacterium* with oral cancer metastasis and aggravated rheumatoid arthritis (6–8). However, the causal relationship and the molecular mechanisms involved are still lacking. While the association between *F. nucleatum* and diseases, including its potential role in pathogenesis, has been recognized, the specific virulence factors driving these processes remain largely unknown. One significant impediment to elucidating the causal relationship between *F. nucleatum* and diseases is the limited availability of efficient genetic tools for comprehensive studies. The genetic manipulability of *F. nucleatum* is limited to only a few reported strains (9–11). Efforts to develop efficient genetic tools for *F. nucleatum* have been impeded by the low transformation efficiency for most strains of this bacterium, and tunable genetic elements at both the transcriptional and translational levels.

Electroporation and conjugation are two primary methods for DNA delivery in bacteria. The efficiency of these transformations is influenced by various factors, including the methods for the preparation of competence cells, electroporation parameters, and the ratio of donor to recipient cells in conjugation assay. One intrinsic factor is the host bacteria’s defense system against foreign DNA. Many bacteria possess large number of innate and acquired defense systems to degrade foreign DNA, which include Restriction-Modification (RM) systems, CRISPR-Cas, abortive infection, prokaryotic Argonaute, and the recently unveiled BREX system (12–15). Of these, restriction-modification systems predominate, present in nearly 95% of all sequenced bacterial genomes, and are commonly considered the greatest barriers to DNA transfer (16). The presence of RM systems was also found to be one of the main reasons for the low transformation efficiency of *F. nucleatum* (17). Careful analysis of recognition sites and their corresponding methyltransferases is essential for developing strategies to enhance transformation efficiency by circumventing RM systems. This can include modifying donor DNA to eliminate recognition sites or engineering intermediary strains to methylate donor DNA before transformation.

The RM system comprises a methyltransferase, which possesses the capability to methylate DNA with host-specificity motifs, and an endonuclease, which could cleave non-methylated DNA (18). These two components show different function and limited conservation in close species, resulting in various diversities of RM systems among related taxa (19). The RM systems comprise four types, Type I-IV. Type I is the most complex one, consisting of three subunits: R, M, and S, encoded by genes located adjacent to the genome, named respectively: *hsdR*, *hsdM*, and *hsdS* (20). An R_2_M_2_S complex is formed to serve functions of methyltransferase and endonuclease (21). The Type II and III RM systems consist of R and M subunits. Type I-III RM systems are characterized by distinct recognition motifs, typically spanning 2 to 9 base pairs. The Type IV RM system is distinct from the modification-blocked Type I-III, which comprises modification-dependent enzymes cutting modified DNA sequence (22). Several approaches have proven effective in enhancing bacterial transformation by overcoming restriction-modification systems, including heat shocks which decreases the activity of endonuclease in the host (23), codon optimization to avoid the RM recognition motifs in the donor DNA (20), removal of the encoded restriction enzymes from the bacterial genome of the host and so on. One of the most efficient way is utilization of an intermediate host with modified methylation patterns to match those of the final host (24). A study used *E. coli* expressing Type II methytransferases as an intermediate host to mimic the DNA methylation patterns of the final host achieved 10^6^-fold increase of the transformation efficiency (25). The previously mentioned methods have been demonstrated to be effective in numerous bacterial species. However, their systematic application to *F. nucleatum* has been hindered by a lack of information regarding the active restriction-modification (RM) system in this species. As sequencing technology advances, PacBio SMRT sequencing is able to accurately identify methylation sites and types (26). This development simplifies and streamlines the application of the strategy to all bacterial strains with a constrained modification system. Previous research used PacBio SMRT sequencing to study the RM system in *Bifidobacterium subspecies* CNCM I-2494, and successfully expressed methyltransferases in *Escherichia coli* that facilitate genetic manipulation of *Bifidobacterium subspecies* (27). Moreover, this approach has the potential to be extended to a wide range of non-model bacteria, including *F. nucleatum* (28). A recent study performed bioinformatic analysis of RM systems in *Fusobacterium* and predicted RM systems with various numbers in all the tested strains (17). Through the expression of only Type II and III methyltransferases in the intermediate host, they achieved a 50-fold increase in the transformation efficiency of *F. nucleatum* (17), highlighting that bypassing the RM system is crucial for enhancing genetic tractability.

In this study, we used PacBio SMRT sequencing and identified the methylated motifs present in two strains of *F. nucleatum*, ATCC10953 and ATCC25586, thereby shedding light on the intricate epigenetic landscape of this pathogen. Subsequently, we identified the methyltransferases responsible for the DNA methylation motifs. Building upon this knowledge, we developed a pipeline aimed at enhancing transformation efficiency in *F. nucleatum* by circumventing restriction-modification (RM) systems, thus opening avenues for more effective genetic manipulation in this species. Furthermore, we introduced a pioneering conjugation strategy tailored to *F. nucleatum*, marking the first application of such a technique in this bacterium. Additionally, we developed a dual-plasmid system, providing a versatile platform for future genetic engineering endeavors, including the implementation of dual-plasmid CRISPR editing strategies. This study not only expands our understanding of *F. nucleatum* biology but also lays the groundwork for future studies aiming to dissect its pathogenic mechanisms.

## Results

### Identification of active Restriction Modification systems by SMRT sequencing

RM systems were shown to be major barriers for DNA transformation in various bacteria. Identification of the recognition sites and their corresponding enzymes are the key to overcome this barrier. The Restriction Enzyme Database (REBASE) (http://rebase.neb.com/cgi-bin/pacbiolist) predicted that both *F. nucleatum* ATCC25586 and ATCC10953 harbors multiple R-M systems, but the active enzymes and their recognition sites are not clear. To identify the active RMs and their recognition motifs, we performed PacBio Single Molecule real-time (SMRT) sequencing to detect DNA methylations on a genome-wide scale and analyzed all base modifications with the two *F. nucleatum* strains (Fig. 1A).

**Fig 1.**
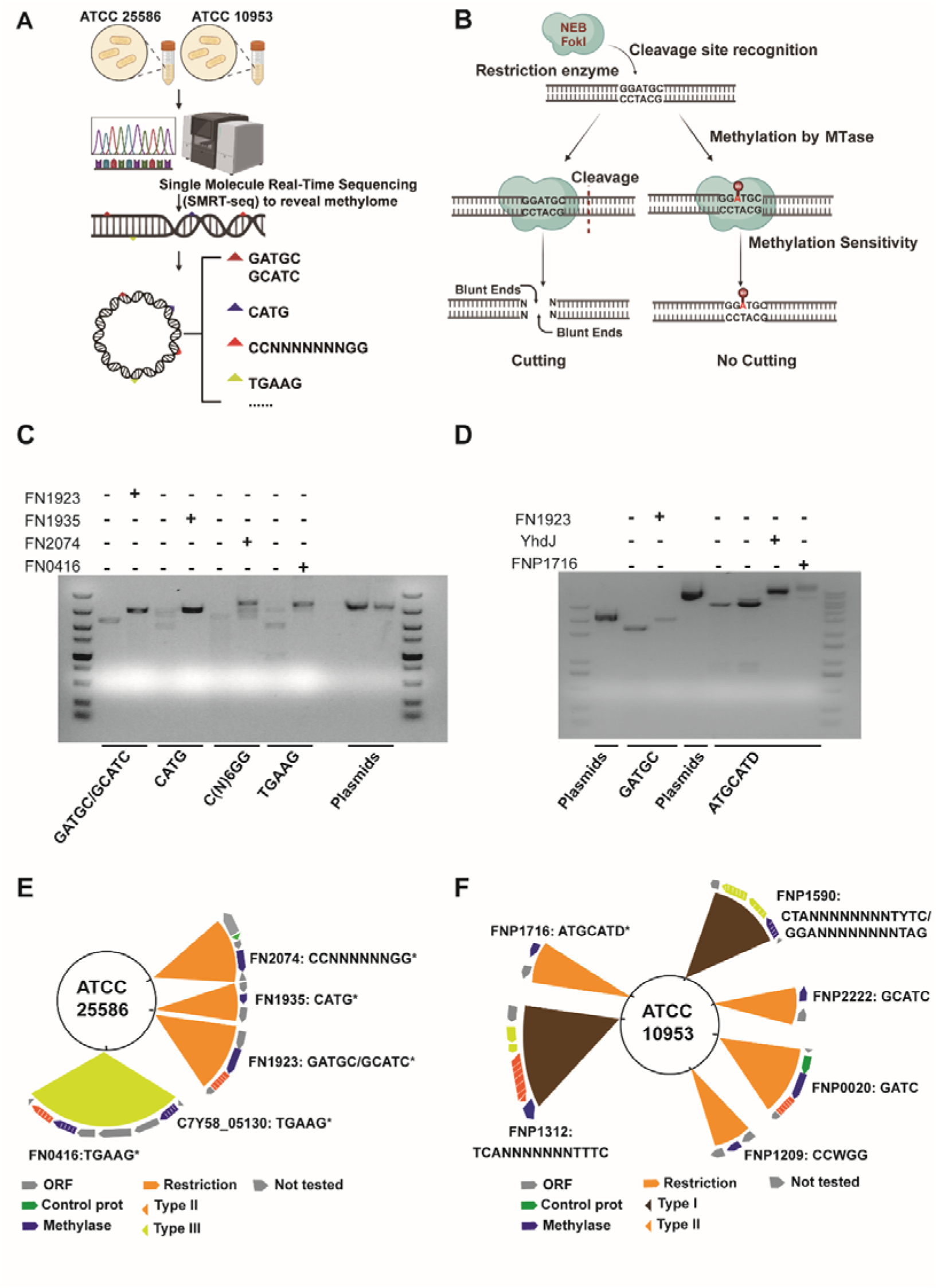
Identification and classification of functional restriction-modification systems and recognition motifs from *F. nucleatum*. **(A)** The pipeline of PacBio SMRT sequencing to identify functional RM systems in *F. nucleatum* ATCC25586 and ATCC10953. **(B-D)** Identify the corresponding methyltransferase for the methylated DNA motifs identified by PacBio Sequencing. **(B)** Determine the activity of DNA methyltransferases (MTases) by the methylation sensitive endonuclease digestion assay. **(C, D)** Identification of the MTases responsible for the methylated DNA motifs. MTases were expressed individually in the methyltransferase-deficient strain *E. coli* ER2976. The *E. coli* ER2976 expressing different MTases were used as an intermediate host for the plasmids. Plasmid pKH9-RM was used for the restriction digestion assay in panel **C** and lanes 1-3 of panel D, whereas plasmid pFN1-AsCas12f was used for lanes 4-8 of panel **D**. “-” and “+” means without or with expression of the MTase in the host *E. coli* ER2976. The tested DNA motifs were labelled at the bottom of figure. **(C)**. Lanes 1-2 shows the identification of MTase for motif GATGC; lanes 3-4 for motif CATG; lanes 5-6 for motif CCNNNNNNNGG; lanes 7-8 for motif TGAAG; lanes 10-11 showed the plasmid pKH9-RM that was not digested by methylation sensitive restriction enzymes. **(D)**. Lanes 1-3 shows the identification of MTase for motif GATGC. Note that lane1 showed the not digested plasmid pKH9-RM as a negative control. Lanes 4-8 were for identification of MTase for motif ATGCATD. Lane4 was negative control of not digested plasmid pFN1-AsCas12f. Note that YhdJ is the MTase reported to methylate ATGCATD motif in *E. coli* and serves as a positive control here. (**E-F**) The summary of the methylated DNA motifs identified by PacBio sequencing and their corresponding MTase. * means the MTase recognition motif was confirmed by the methylation sensitive endonuclease digestion assay as presented in panels **C** and **D**. Motifs identified solely through bioinformatic analysis are not marked with *.

Two types of DNA modifications, including N6-methyladenine (6-mA) and N4-methylcytosine (4-mC), were detected by SMRT sequencing in *F. nucleatum* ATCC25586, including 8 different motifs evenly distributed over the chromosome (Table 1, Figure S1A). However, in *F. nucleatum* ATCC10953, only 6-mA were detected, including 5 motifs across the chromosome and plasmid (Table 2, Figure S1BC). Notably, our analysis unveiled one previously unreported DNA methylation motif “TGAAG” in *F. nucleatum* ATCC25586, expanding the diversity of MTases in bacteria. The results showed that sequence of DNA methylation motifs between the two strains are significantly different. One difference is that ATCC10953 harbors 5’-G**A**TC-3’ 6-mA modifications, whereas ATCC25586 doesn’t. This explains the observation that plasmid isolated from ATCC10953 cannot be successfully transformed into ATCC25586 (17). Interestingly, *F. nucleatum* ATCC10953 harbors a plasmid pFN3, which is 11934 bp (NCBI Reference Sequence: NC_009506.1). The SMRT sequencing results showed that the RM recognition motifs on the plasmid were all methylated. Sequence search showed that this plasmid has only one GATC motif, much less frequent than the genome of ATCC10953 (1513 motifs of 2.17 Mbp), or native plasmids in other *F. nucleatum* strains, like pFN1 (2 GATC motifs of 5887 bp) or pKH9 (2 GATC motifs of 4975 bp), indicating methylation of GATC plays a role in acquisition of exogenous DNA for *F. nucleatum* ATCC10953.

**Table 1.**
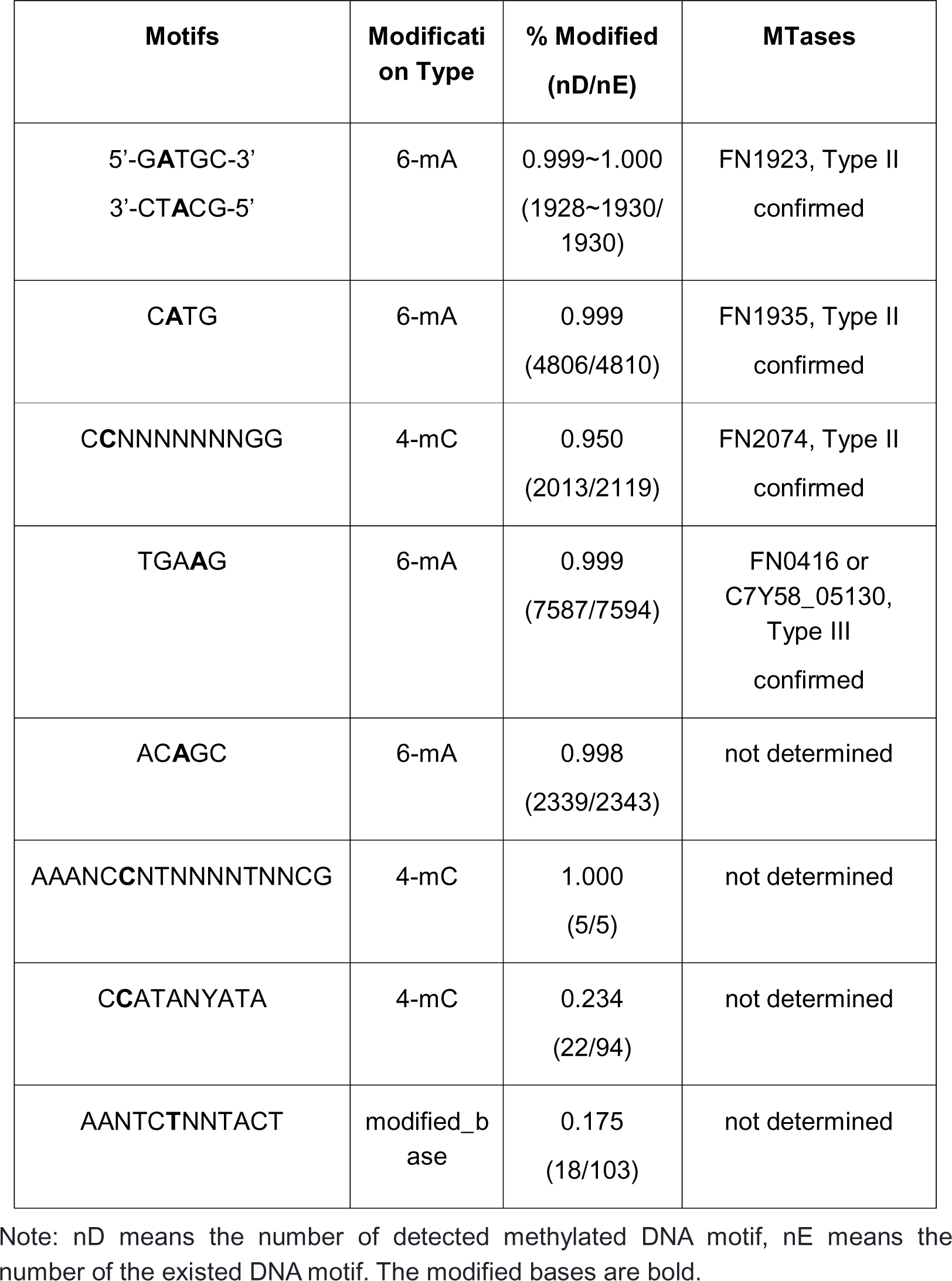
DNA Methylation motifs detected with *F. nucleatum* ATCC25586.

**Table 2.**
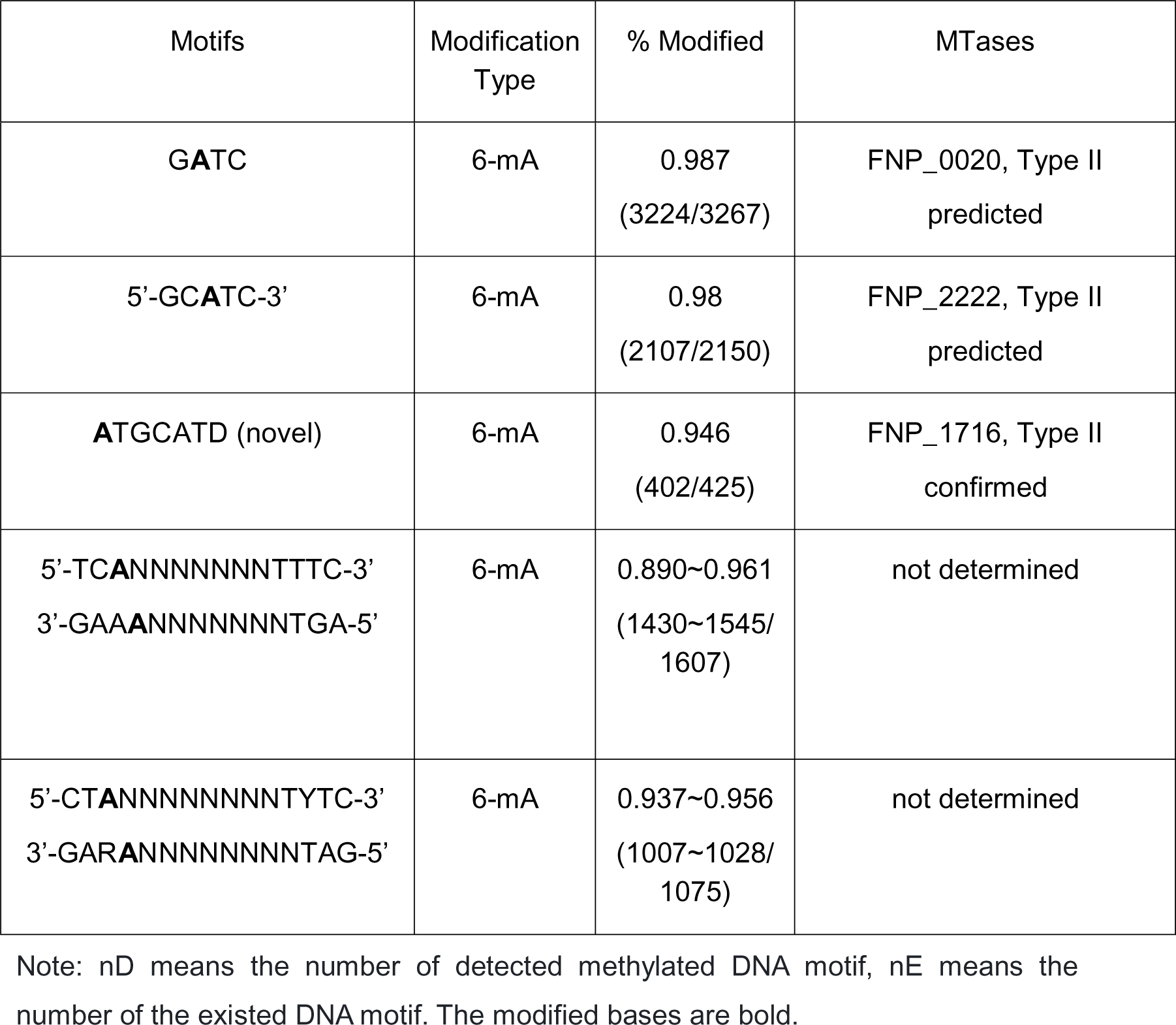
DNA Methylation motifs detected with *F. nucleatum* ATCC10953 chromosome.

### Characterization of the methyltransferase for the identified DNA methylation motifs

To pinpoint the methyltransferases (MTases) responsible for the DNA methylation motifs, we devised an assay utilizing established methylation-sensitive endonucleases to differentiate between methylated and unmethylated DNA motifs (Fig. 1B). We cloned and expressed putative MTases from *F. nucleatum* ATCC25586 and ATCC10953 in *E. coli* ER2796, a strain lacking endogenous MTases (29), yielding a set of intermediary *E. coli* hosts, each expressing a distinct MTase. Subsequently, plasmids containing a combination of the tested DNA motifs and endonuclease recognition sites were introduced into these intermediary *E. coli* hosts. The methylation status of the DNA motifs on the plasmid was then assessed using the assay shown in Fig. 1B.

We mainly focused on strain *F. nucleatum* ATCC25586 and identified the MTases for 4 DNA motifs in this strain. The results showed that FN1923 (GenBank: AAL94022.1) was responsible for the N6-methyladenine of (5’-GATGC-3’/3’-CTACG-5’) motif; FN1935 (GenBank: AAL94034.1) for the 6-mA of 5’-CATG-3’; FN2074 for 5’-C(N)_6_GG-3’; FN0416 (GenBank: AAL94619.1) for 5’-TGAAG-3’ (Fig. 1CE). The MTase responsible for the DNA motif 5’-ACAGC-3’ was not identified through the predicted RM systems listed in the Rebase database. However, it has been reported that this motif can be methylated by known MTases found in other bacteria, as annotated in the Rebase database, suggesting that these MTases could be leveraged to provide this modification. For *F. nucleatum* ATCC10953, we found that FNP_1716 (GenBank: EDK89489.1) is responsible for the methylation of 5’-ATGCATD-3’, which shared the same methylation motif of YhdJ from *E. coli* DH5α (Fig. 1DF). It’s interesting to notice that the DNA motif (5’-GATGC-3’/3’-CTACG-5’) was fully methylated in ATCC25586 by FN1923, but in ATCC10953 it is hemi-methylated with only the strand with 5’-GCATG-3’ modified. The identified and predicted MTases for each DNA methylation motifs were summarized in Table 1 and Table 2.

### GATC 6-mA modifications and its role in transformation of *F. nucleatum*

It’s well known that Dam of *E. coli* can perform 6-mA of 5’-GATC-3’ motif (30), which was identified in *F. nucleatum* ATCC10953, but not in *F. nucleatum* ATCC25586. We hypothesized that *F. nucleatum* ATCC10953 has a homolog of Dam, which is absence in ATCC25586. Protein domain analysis identified that FNP_0020 (GenBank: EDK87839.1) of *F. nucleatum* ATCC10953 harbors a Dam domain (Fig. 2A), indicating its role in 6-mA of 5’-GATC-3’ motif. Expression of FN_0020 in E. coli ER2796 effectively modifies GATC sites on plasmids, making them susceptible to DpnI digestion (Fig. 2B). To understand the conserved domains about the FNP_0020 protein, a systematic inquiry was conducted in the NCBI Conserved Domain Database (CDD). Eight MTases previously reported to methylate the 5’-GATC-3’ motif with 6-mA were selected and aligned with FNP_0020. This alignment revealed 16 conserved amino acid residues within the 18-265 amino acid residue domain (Fig. 2A).

**Fig 2.**
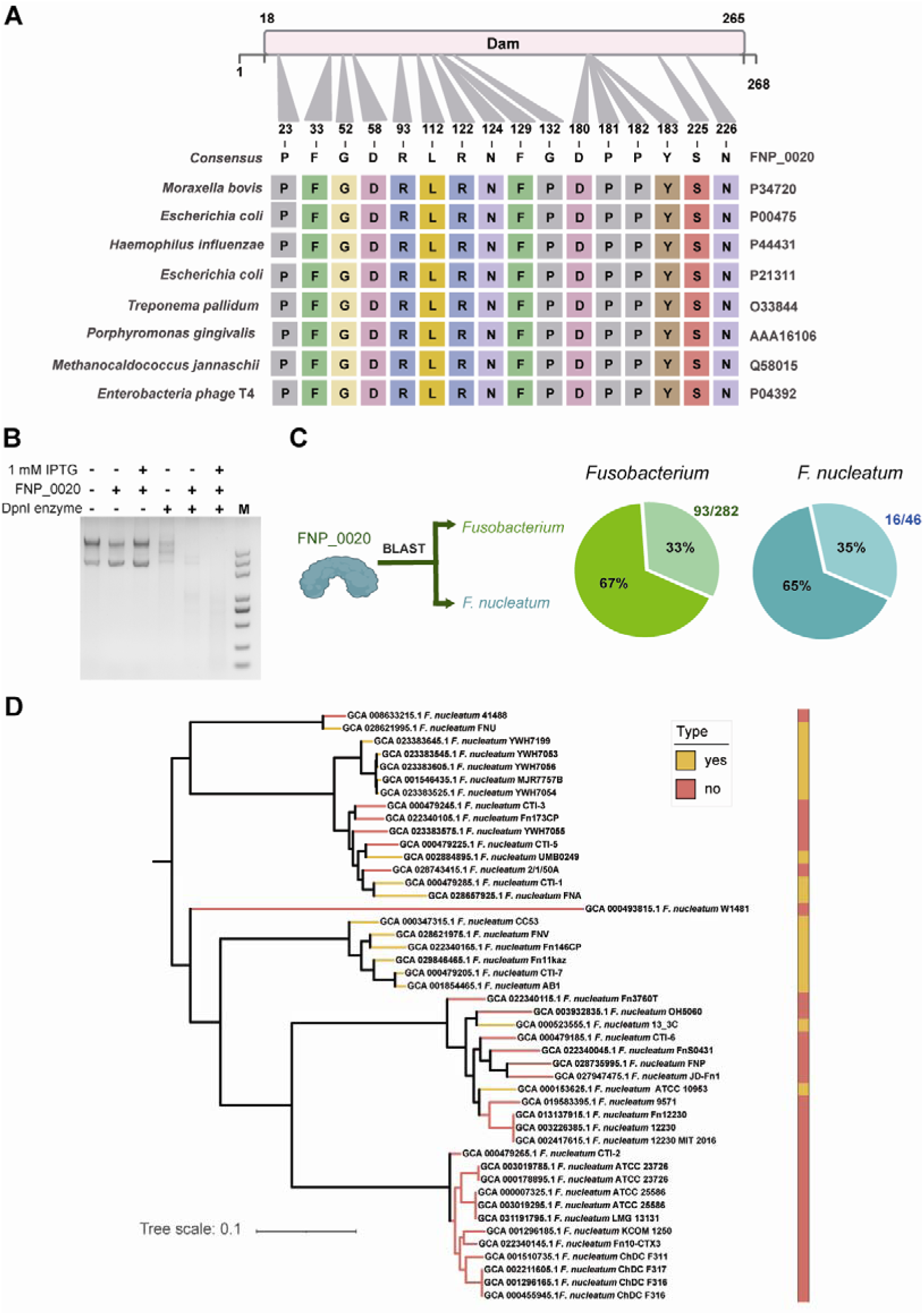
Distribution of FNP_0020 methylating GATC motif in *Fusobacterium* genus and *F. nucleatum* species identified by bioinformatic analysis. (**A**) Multiple alignment of the amino acid sequence of FNP_0020 with eight MTases methylating GATC in other species. The result showed that a highly conserved region ranging from amino acids 18 to 265 of the CARD Dam domain. (**B**) Identification of FN_0020 as the MTase for G**A**TC (6-mA) mofidication. FN_0020 was expressed in the host *E. coli* strain under IPTG- inducible promoter. DpnI digestion was used to distinguish the G**A**TC modifications on the plasmids. (**C**) Distribution of FNP_0020 proteins in *Fusobacterium* genus and *F. nucleatum*. The pie chart shows the percentage of strains harboring a homolog of FNP_0020 methyltransferase. The lighter shades represent the percentages of strains with homolog, whereas the darker shades indicate the absence of homolog. (**D**) The phylogenetic tree of 46 *F. nucleatum* strains accompanied by the distribution of homologs of FNP_0020. Orange indicates the presence of a homology of FNP_0020 in the strain, whereas red indicates absence.

The methylation pattern of 5’-GATC-3’ of the target strain is a major factor for choosing the right intermediary *E. coli* host to provide matched donor DNAs (31). To test the impact of GATC methylation on transformation efficiency of *F. nucleatum*, we isolated the shuttle vectors from *E. coli* DH5α (Dam+) and *E. coli* ER2796 (Dam−), and then electro-transformed the GATC methylated or unmethylated vectors into ATCC10953 and ATCC25586. We observed successful transformation of GATC methylated shuttle vectors into ATCC10953, while unmethylated vectors failed to transform. For ATCC25586, 5-10 colonies were obtained when transform 3 microgram unmethylated shuttle vectors, while no colony appeared with methylated vectors. The low transformation in ATCC25586 is due to the presence of other recognition motifs on the vector. The above observations confirmed that the right methylation of GATC is crucial for transformation in *F. nucleatum*

To provide guide information to choose the right intermediary *E. coli* host for different *F. nucleatum* strains, we analyzed the distribution of FNP_0020 in *Fusobacterium* genus and *F. nucleatum*, respectively. In total, 282 genome datasets of the *Fusobacterium* genus were procured from the NCBI database. Within these datasets, 93 genomes were identified to contain a homolog of FNP_0020 protein, accounting for approximately 33% of the total (Fig. 2B). Concurrently, 46 genomic datasets of *F. nucleatum* were acquired and analyzed. In the set of 46 downloaded genomic sequences, approximately 35% exhibit similarity to the FNP_0020 protein, accounting for 16 sequences in total (Fig. 2B). The Phylogenetic tree was made to show the distribution of FNP_0020 homolog in *F. nucleatum* (Fig. 2C). It is noteworthy that the genome of *F. nucleatum* ATCC25586 does not have the homologous protein, in consistent with the SMRT sequencing result.

### Bypassing the restriction barrier enhances transformation efficiency in *F. nucleatum*

Based on the identified DNA methylation motifs and their corresponding MTases, we next developed an effective pipeline to enhance the transformation efficiency in *F. nucleatum* (Fig. 3A). This pipeline involved expression of a selected subset of MTases from the *F. nucleatum* in an intermediary *E. coli* host (Fig. 3A). Two *E. coli* strains, ER2796 and DH5α, were selected as the background strains for inserting the MTase genes to engineer the intermediator (Fig. 3B). *E. coli* DH5α, which is Dam positive for 6- mA 5’-GATC-3’, was used as the intermediary *E. coli* host for *F. nucleatum* ATCC10953, which contains 6-mA 5’-GATC-3’ motif. Meanwhile, *E. coli* ER2796, a MTase-free strain, was utilized to express MTases for constructing the intermediary *E coli* host for *F. nucleatum* ATCC25586. To engineer the intermediary *E. coli* host for *F. nucleatum* ATCC25586, the genes of three MTases—FN2074, FN1935, and FN1923—were inserted into the chromosome of *E. coli* ER2796 via tri-parental mating (Fig. 3B). Similarly, to engineer the intermediary *E. coli* host for *F. nucleatum* ATCC10953, the genes of two MTases—FN1923 and FNP_1716—were inserted into the chromosome of *E. coli* DH5α using tri-parental mating (Fig. 3B). To assess the effectiveness of this pipeline, we compared the transformation efficiency of plasmids with or without methylation using the intermediary *E. coli* hosts. The plasmid pFN1-catP, which contains multiple RM recognition sites of *F. nucleatum* ATCC25586 (Fig. 3C), was utilized for testing *F. nucleatum* ATCC25586. The transformation results showed that pFN1-catP isolated from empty *E. coli* ER2796 produced no transformants. However, the same plasmid isolated from *E. coli* ER2796 expressing the three MTases was efficiently transformed, yielding approximately 5000 CFU per microgram of plasmid (Fig. 3D). Similar results were obtained with the transformation of pFN1-P_4.5s_-NanoLuc into *F. nucleatum* ATCC10953 (Fig. 3EF). It’s noticeable that the sole 6-mA of 5’-GATC-3’ significantly improves the efficiency of transformation with *F. nucleatum* ATCC10953. These findings support the notion that bypass the RMs via the pipeline described in Fig. 3A can significantly enhance the electro transformation efficiency of *F. nucleatum*, reaching up to 10^3^-fold.

**Fig 3.**
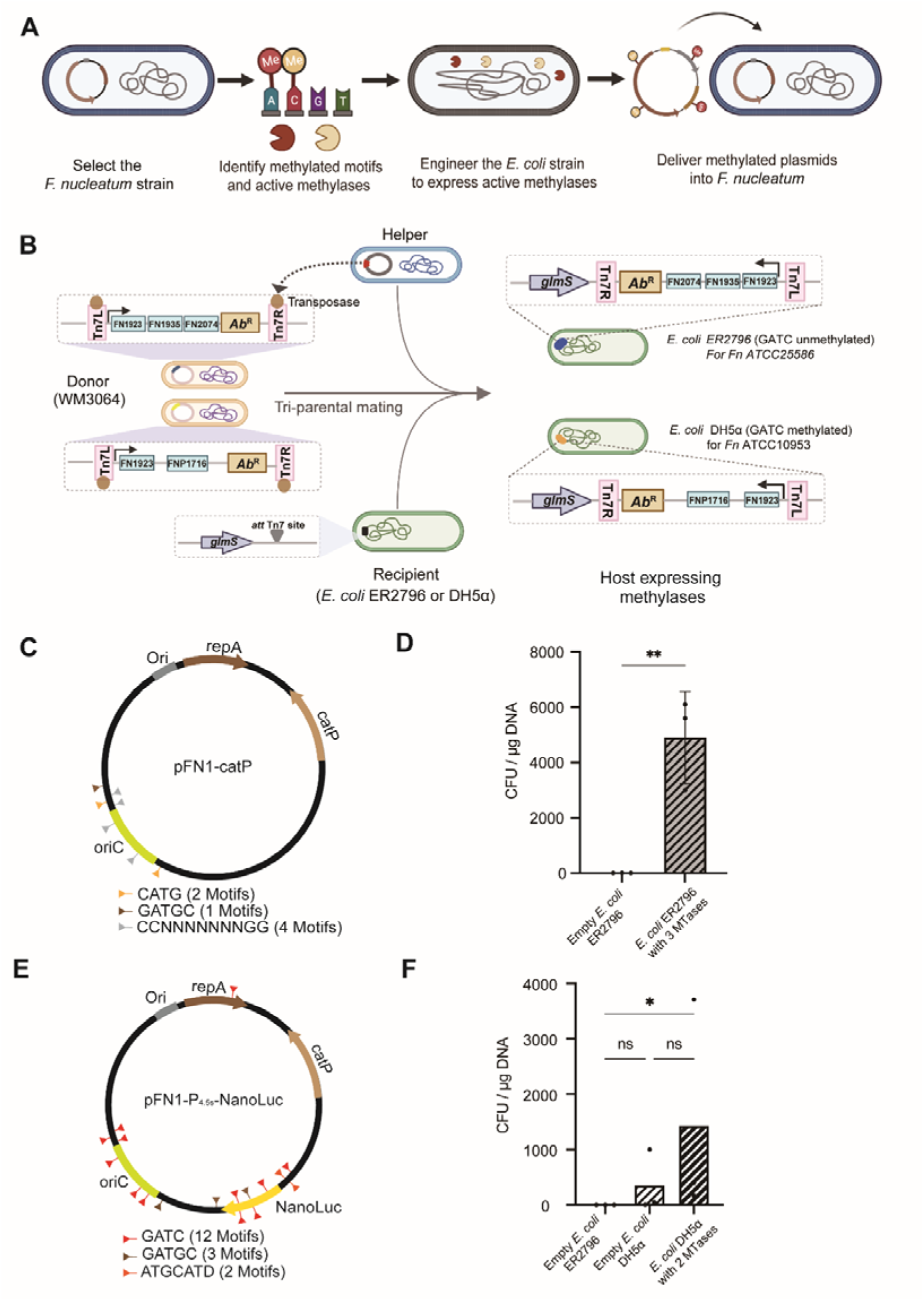
Transformation efficiency of *F. nucleatum* ATCC 10953 and ATCC 25586 with various shuttle plasmids. **(A)** The systematic pipeline for enhancement of transformation efficiency of *F. nucleatum* bypassing restriction-modification systems. **(B)** Construction of the intermediator host *E. coli* strains to mimic methylation patterns of *F. nucleatum* ATCC25586 and ATCC10953. Tri-parental conjugation and Tn7 transposition was used to integrate the expression cassette of multiple MTases onto *E. coli* genome. (**C**-**D**) Vector map of shuttle plasmid pFN1-catP and transformation efficiency tested with this vector in *F. nucleatum* ATCC25586. Transformation efficiency of *F. nucleatum* ATCC25586 strain with shuttle plasmids prepared from *E. coli* ER2796 expressing MTases. **(E-F**) Vector map of shuttle plasmid pFN1-P_4.5s_- NanoLuc and transformation tested with this vector in *F. nucleatum* ATCC10953. **(D)** Transformation efficiencies shown are averages of at least three replicates ± SD. Unpaired t test was used to calculate p values in (D). * p < 0.05, ** p < 0.01, *** p < 0.001. one-way ANOVA was used to calculate p values in (F). * p < 0.05, ** p < 0.01, *** p < 0.001.

### Efficient delivery of plasmids into *F. nucleatum* by conjugation

Our previous results indicated that restriction-modification systems (RMs) are significant barriers to DNA transformation, and bypassing these RMs can significantly enhance electro-transformation efficiency. It has been reported that conjugation machinery transfers single-stranded DNA (ssDNA), which can evade restriction enzyme cleavage until it becomes properly methylated (32). This mechanism may offer an advantage over electroporation, which utilizes improperly methylated DNA (33). While conjugational transformation of DNA has been widely employed in various bacteria, this strategy has not yet been reported for *F. nucleatum*. Therefore, we made efforts to establish a method for introducing shuttle vectors into *F. nucleatum* through bi-conjugation. The conjugation donor strain *Escherichia coli* WM3064 was employed, which has RP4 *tra* genes integrated on the chromosome and is auxotrophic for diaminopimelic acid enabling removal of this strain after conjugation (34). The pFN1- based shuttle vector was equipped with the original site of transfer, known as the oriT site, to confer mobility (35) (Fig. 4A). To set up appropriate conditions for conjugation, we evaluated the aerotolerance of *F. nucleatum* and observed a significant decrease in viability after one hour air exposure. Therefore, we conducted the conjugation process under anaerobic conditions. The *E. coli*-*F. nucleatum* bi-conjugation transformation was performed for *F. nucleatum* ATCC10953 as described at the materials and methods section, and we successfully obtained thiamphenicol-resistant conjugants. The presence of pFN1 in the transconjugants was further confirmed by PCR (Fig. 3C). We further optimized the conjugation procedure to improve efficiency by adjusting the donor-to-recipient ratio. Our results demonstrated that increasing the donor-to-recipient ratio enhances conjugation efficiency (Fig. 4D). In summary, through the design and optimization of conjugation procedures, we have developed an anaerobic conjugation-based system for *F. nucleatum*. It’s worth to mention that the conjugation enables efficient transfer of DNA into *F. nuceatum* without the need of MTase methylation. The conjugation DNA delivery strategy is expected to overcome the genetic intractability caused by the RM system barriers associated with *F. nucleatum*.

**Fig 4.**
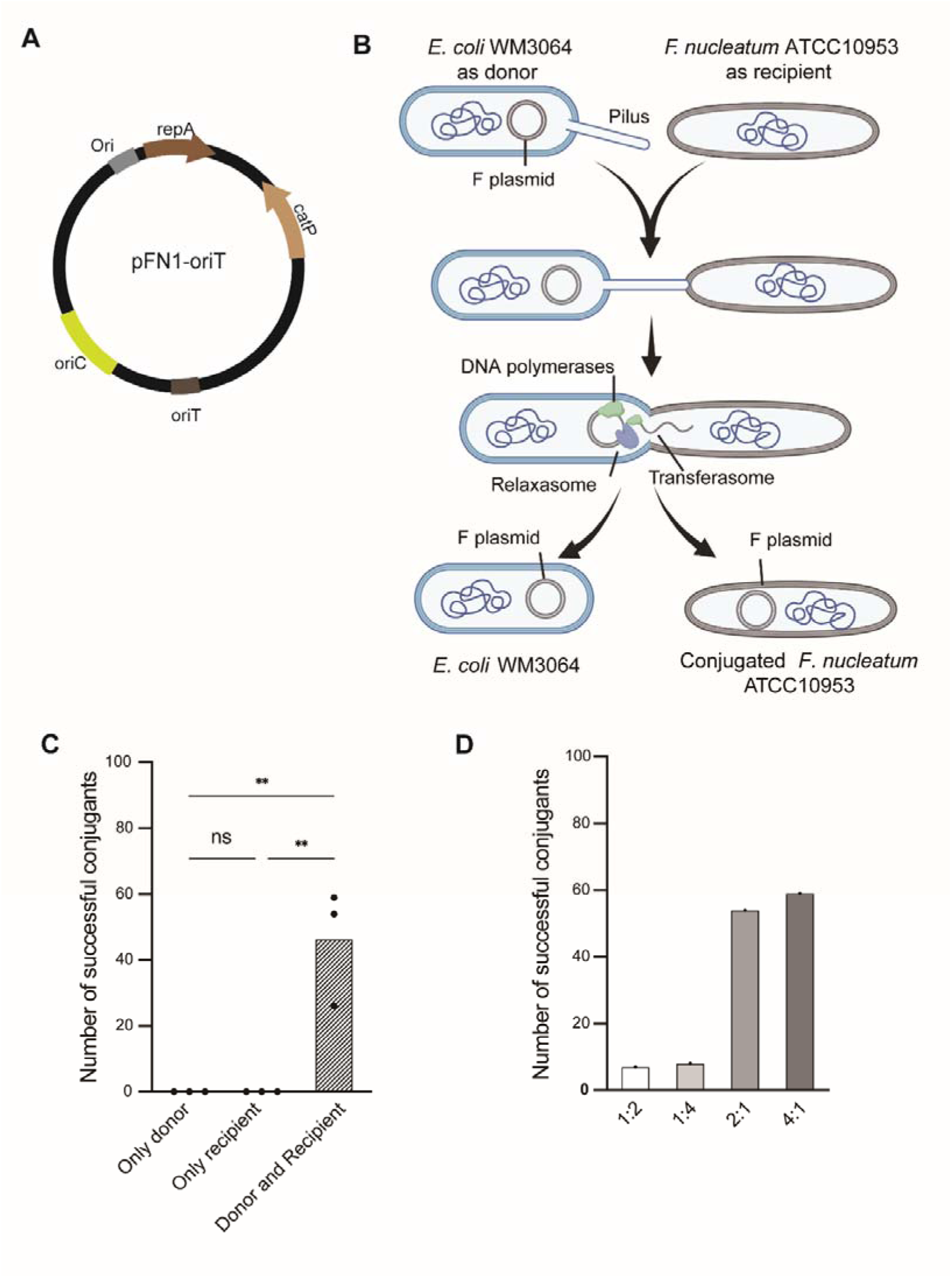
A conjugational transfer method for plasmid delivery in *F. nucleatum*. **(A)** The Schematic map of the pFN1-oriT shuttle plasmid enabling conjugation transformation for *F. nucleatum* - *E. coli* is shown. Note that an oriT site has been added to enable conjugational transformation. **(B)** An overview of bi-conjugation transformation between *F. nucleatum* and *E. coli* is provided*. E. coli* WM3064 with pFN1-oriT was used as the donor strain, and *F. nucleatum* ATCC10953 was the recipient strain. **(C)** The Conjugation transformation efficiency tested with *F. nucleatum* ATCC10953. “Only donor” and “Only recipient” were set as negative control. Conjugant group is the one with both donor and recipient for successful conjugation. The procedure for conjugation was descripted in detail in the methods section. **(C)** The conjugation transformation efficiency at different donor-to-recipient ratios was evaluated, and one representative result is presented here out of two repeated experiments.

### Quantitative analysis of various promoters with different strength for gene expressions

With the improved transformation efficiency achieved through the pipeline to bypass RM systems, we are now able to transform and test diverse genetic elements in *F. nucleatum*. This success has encouraged us to explore a series of promoters, which are key genetic elements, for gene expression with varying strengths in *F. nucleatum*. To facilitate the evaluation of promoter strength, we first constructed a luciferase reporter system in *F. nucleatum*. We chose Nano-Luciferase, a novel bioluminescence platform with enhanced stability, smaller size, and over a 150-fold increase in luminescence compared to established systems (36), for constructing the reporter system (Fig. 5A). Subsequently, we selected and cloned a series of promoter to drive the *nano-luc*, and performed luciferase assay to evaluate the strength of the promoters (Fig. 5B). In total 8 promoters were tested, including the promoter of 4.5s rRNA of *F. nucleatumn* (P_4.5s_); the promoter of *fomA* gene of *F. nucleatum* (P_fomA_) ; the promoter of *fdx* gene of *Clostridium sporegenes* (P_fdx_) (37); the promoter of pGM-ACBQ plasmid used for *F. nucleatum* (P_pmhl_) (38); the promoter of *ermG* from pMT494 plasmid used for *Bacteroides thetaiotaomicron* (P_MT494_) (39); the promoter of pGM-ACBQ plasmid used for *F. nucleatum* (P_thl_) (38); a xylose-inducible promoter and riboswitch combination system used for *F. nucleatum* (P_xylose_) (40), the anhydrotetracycline-inducible promoter from pRPF185 plasmid used for *Clostridium difficile* (P_tetR_) (41). The promoters were tested in both *F. nucleatum* ATCC25586 and ATCC10953, and the result showed that the promoter strength was generally consistent between the two strains, with some exceptions. For example, P_pmhl_ was much stronger in ATCC10953, whereas P_MT494_ is much stronger in ATCC25586. Our results showed that the fomA promoter (P_fomA_) is one of the strongest promoters in both strains, which is consistent with the RNA-seq result performed in multiple *F. nucleatum* strains (42). As expected, the uninduced P_xylose_ and P_tetR_ showed the lowest activity. The tested promoters displayed a wide range of strengths, with luciferase activity ranging from 10^3^ to 10^7^. We anticipate that the promoters tested here can serve as prototypes for the development of promoters with varying strengths.

**Fig 5.**
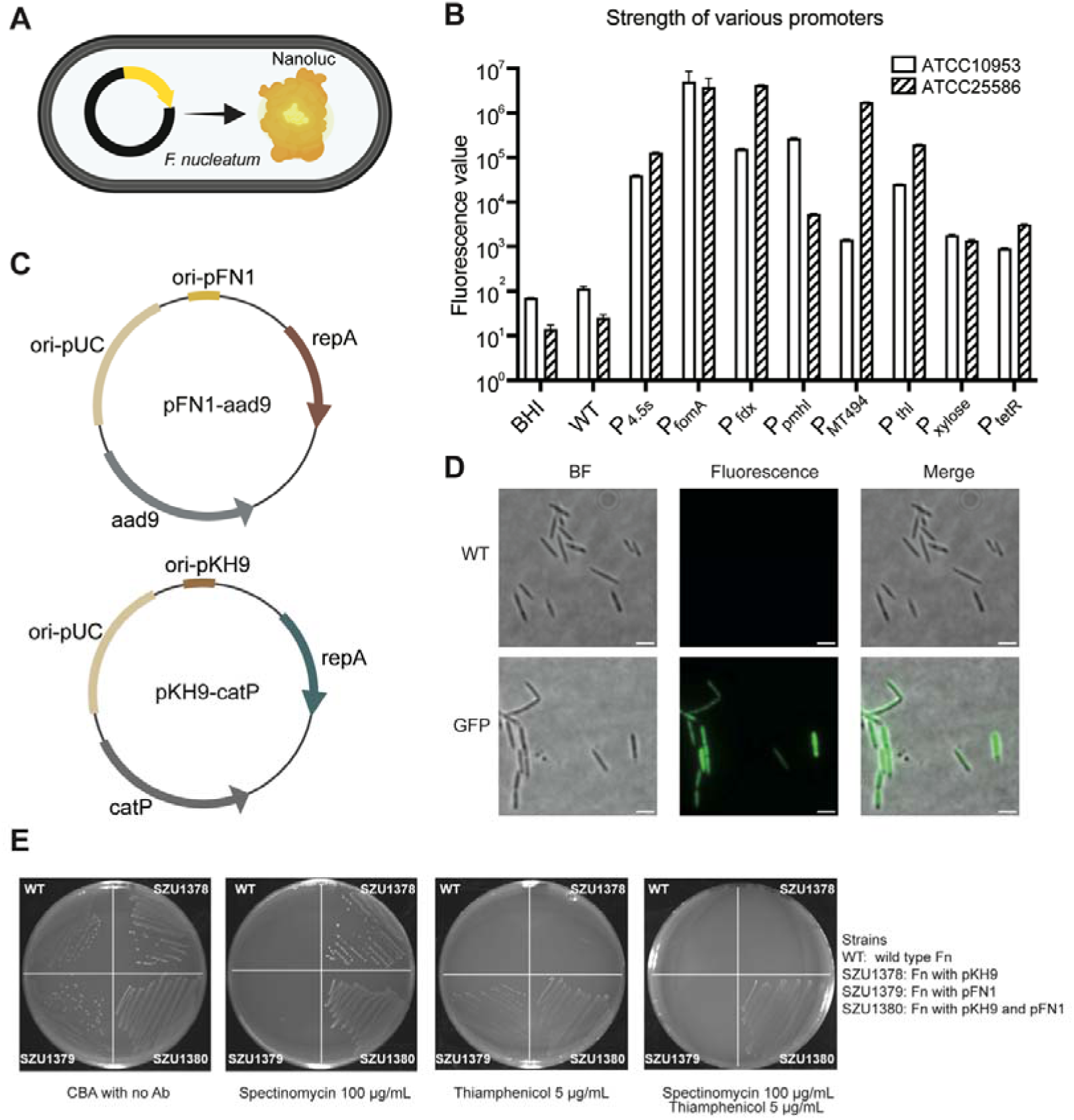
Genetic manipulation elements for *F. nucleatum*. **(A)** The NanoLuc-based luciferase reporter system in *F. nucleatum* is shown. **(B)** Quantification of the strength of various promoters with the luciferase reporter system in both *F. nucleatum* ATCC10953 and ATCC25586 strains. BHI represents the medium as a negative control, WT represents the wild type strain without the vector for expression of NanoLuc. P_4.5s_ represents the promoter of 4.5s rRNA of *F. nucleatum*; P_fomA_ is the promoter of *fomA* gene of *F. nucleatum*; P_fdx_ is the promoter of *fdx* gene of *Clostridium sporegenes* (37); P_pmhl_ is the promoter of pGM-ACBQ plasmid used for *F. nucleatum* (38); P_MT494_ is the promoter of ErmG from pMT494 plasmid used for *Bacteroides thetaiotaomicron* (39); P_thl_ is the promoter of pGM-ACBQ plasmid used for *F. nucleatum* (38); P_xylose_ is a xylose-inducible promoter and riboswitch combination system used for *F. nucleatum* (40); P_tetR_ is the anhydrotetracycline-induce promoter of pRPF185 plasmid used for *Clostridium difficile* (41). **(C)** Maps of the two identified compatible shuttle vectors, pFN1-catP and pKH9-aad9 for *F. nucleatum*. Vector pFN1-catP: ori-pUC is the original site for replication in *E. coli*; ori-pFN1 and *repA* is for replication in *F. nucleatum*; *aad9*, encoding spectinomycin adenyltransferase, is the resistant selection marker. Vector pKH9-aad9: ori-pUC is for replication in *E. coli*; ori-pKH9 and *repA* are for replication in *F. nucleatum*; *catP*, encodind the chloriamphenicol/thiamphenicol acetyltransferase, is the resistant selection marker. **(D)** Exogenous gene expression in *F. nucleatum* with the shuttle vector pFN1. A gene cassette with *gfp*, encoding the green fluorescent protein, driven by a constitutive promoter was cloned into pFN1. The recombinant pFN1-gfp was then transformed into *F. nucleatum* ATCC25586. WT means the wild type *F. nucleatum* ATCC25586; GFP means *F. nucleatum* ATCC25586 transformed with the pFN1-gfp plasmid. Microscopy was performed with phase contrast, green fluorescent channel. The merged images of both phase contrast and green fluorescent channel were also shown. The scale bars are 2 μm. (**E**) Compatibility of the pKH9 and pFN1 in *F. nucleatum*. Growth test of transformants with pFN1 (SZU1378), pKH9 (SZU1379) or both (SZU1380). Wild type *F. nucleatum* was set as control (WT). Growth was tested on Columbia agar plate supplied with 5% sheep blood, with addition of spectinomycin, thiamphenicol or both.

### Establishment of a dual-plasmid system with compatible origins of replication in *F. nucleatum*

To advance the development of genetic editing tools for *F. nucleatum*, we devoted significant effort to establish a dual-plasmid system. The keys for the dual-plasmid system are compatible resistant selection markers and origins of replication. The *catP* gene, encoding chloramphenicol acetyltransferase, has been widely utilized in *F. nucleatum* due to its proven efficiency, making it our initial choice for the antibiotic-resistant marker. Additionally, we explored several other antibiotic resistance genes for both *F. nucleatum* ATCC25586 and ATCC10953. Our investigation revealed that *aad9* (NCBI: WP_063246869.1), encoding spectinomycin adenyltransferase, offers efficient selection in both strains with 100 μg/ml spectinomycin. Streptomycin and kanamycin were also evaluated as potential antibiotic selection markers. However, during our testing, we observed that *F. nucleatum* ATCC10953 exhibited robust growth even in the presence of 100 μg/ml streptomycin, while ATCC25586 generated false positive colonies when exposed to 100 μg/ml kanamycin. As a result, we opted to utilize *aad9* as the second selection marker alongside *catP* for the construction of the dual-plasmid system.

To identify compatible origins of replication, we conducted a search for reported native plasmids in *F. nucleatum*. Our search yielded four plasmids, namely pFN3 from *F. nucleatum* ATCC10953, pFN1 and pPA52 (GenBank: AF022647) from *F. nucleatum* FDC 27–17, pFN1 (GenBank: AF159249.1) from *F. nucleatum* 12230, and pKH9 (GenBank: AF295336.1) with an unknown source strain. Sequence analysis of origin sites and replication proteins revealed a potential similarity between the replication systems of pFN1 and pPA52. As a result, we focused our investigation on the replication systems of pFN1, pFN3, and pKH9. We initially constructed pFN1-catP and pKH9-catP plasmids and verified their successful transformation into *F. nucleatum*. Subsequently, we expressed green fluorescent protein (GFP) in *F. nucleatum* ATCC25586 using the pFN1-catP backbone (Fig. 5D), demonstrating the stable replication capability of the vector in *F. nucleatum*. We subsequently assessed the compatibility of the plasmids with each other. Transformation of pFN1-aad9 and pKH9-catP (Fig. 5C) into *F. nucleatum* ATCC10953, which harbors pFN3, demonstrated that both plasmids could stably replicate alongside pFN3 in ATCC10953. This result indicates that both pFN1-aad9 and pKH9-catP are compatible with pFN3. We subsequently transformed pFN1-aad9 and pKH9-catP separately or together into *F. nucleatum* ATCC25586. The resistance of the wild-type strain, *F. nucleatum* ATCC25586 carrying only pFN1-aad9, only pKH9-catP, or both plasmids was then tested using CBA plates supplemented with different antibiotics. The results demonstrated that transformation of the plasmids conferred corresponding resistance, with pFN1-aad9 and pKH9-catP coexisting in *F. nucleatum*, thereby providing resistance to both thiamphenicol and spectinomycin (Fig. 5E). In summary, we have successfully established a dual-plasmid system for *F. nucleatum*, which should pave the way for the development of efficient genetic editing tools in this recalcitrant bacterium.

## Discussion

Many bacteria possess restriction-modification (RM) systems that recognize specific structural features within DNA sequences and identify foreign DNA (18). Our previous attempts to transform plasmids for gene expression or knockout in *F. nucleatum* consistently failed, and other studies also struggled to delivery DNAs into some strains of this bacterium (6). This prompts us to investigate the presence of RM systems and their role in restriction of foreign DNAs in this species (44). We obtained information indicating the presence of multiple predicted RM systems in *F. nucleatum* from the NEB Rebase database. To validate this information and gather comprehensive data, we conducted functional RM system detection through PacBio SMRT sequencing and subsequent bioinformatic analysis. We found 8 DNA methylation motifs in *F. nucleatum* ATCC25586 and 5 in ATCC10953, indicating multiple active RMs in both strains. Further studies identified the MTases responsible for these modifications. To bypass the major RM systems of *F. nucleatum* for transformation efficiency improvement, we constructed a set of intermediary *E. coli* hosts expressing distinct MTase individually, or combined, to provide plasmids with methylation modifications. We found that solely methylation of the DNA with each MTase couldn’t significantly enhance plasmid transformation efficiency. However, combining three MTases yielded a significant increase in ATCC25586, in consistent with previous research in other bacteria (17). The phenomenon may be attributed to the prevention of cleavage at the potential restriction sites, where even a single cut of the plasmid would likely impede its successful transformation. By employing 3 MTases from *F. nucleatum* ATCC25586 to methylate plasmids, we successfully made a remarkable 1000-fold enhancement in transformation efficiency, reaching 5000 transformants per µg plasmid. Note that we didn’t include the MTase for 5’-ACAGC-3’ methylation, which can be methylated with known MTase from other bacteria (refer to Rebase database). We suspect that further improvement of transformation efficiency of *F. nucleatum* ATCC25586 can be achieved by addition of a MTase for this motif. It is worth mentioning that adopting the MTases modifying the same motif from other bacteria is a good option to engineer the intermediary *E. coli* hosts. For instance, YhdJ MTase in *E. coli* DH5α corresponds to the ‘ATGCATD’ motif.

Our study revealed the presence of 6-mA modification at the 5’-GATC-3’ motif in *F. nucleatum* ATCC10953, but not in *F. nucleatum* ATCC25586. This finding supports previous observations indicating that GATC modification is strain-specific within this species (30). Further studies showed that proper methylation of GATC is essential for successful transformation in both *F. nucleatum* ATCC25586 and *F. nucleatum* ATCC10953. These observations indicate the existence of type IV RM system or DpnI or DpnII homolog systems in *F. nucleatum*, which needs further exploration. Type IV enzymes target methylated DNA (45). For example, Mrr or McrBC, a type IV restriction endonuclease involved in the acceptance of modified foreign DNA, restricts both adenine- and cytosine-methylated DNA, but not 5’-GATC-3’ sequence, and its mechanism is not clear ^[43,44]^. It’s well known that Dam of *E. coli* performs the 6-mA of 5’-GATC-3’. By bioinformatic analysis, we identified a MTase FNP_0020 with dam domain in *F. nucleatum* ATCC10953, while such MTase is lacking in ATCC25586 as expected. We further analyzed the distribution of FNP_0020 in *Fusobacterium* genus and specifically in *F. nucleatum*, and find that approximately 33-35% of the population are with FNP_0020. Based on these observations, we suggest to select the appropriate *E. coli* host (Dam+ or Dam-) to match the GATC methylation pattern of the *F. nulceatum* strains, thereby improving the transformation efficiency. This pipeline should be universally applicable to other *F. nucleatum* strains.

For most of the strains, matching the GATC methylation pattern should be able to provide successful transformation, while matching other methylations pattern should be the keys for high transformation efficiency. The limitation of the pipeline proposed in this study is that it requires the tailored construction of the intermediary *E. coli* host. However, it offers significant advantages over previous methods in terms of improving transformation efficiency, with an increase of up to 10^3^-fold compared to a 50-fold increase reported previously (17). This enhancement is crucial for conducting high-throughput genetic screens with this bacterium. The significantly improved transformation efficiency provides a solid foundation for establishing high-throughput genetic studies with this bacterium, such as constructing transposon and CRISPR interference mutant libraries(47, 48). Considering that the dCas9-based CRISPR interference system has been reported to be applicable to *F. nucleatum* (49), we anticipate that CRISPR interference-based high-throughput genetic screens could provide new insights into the pathogenesis of this bacterium.

To establish genetic tools for *F. nucleatum*, several key considerations and challenges need to be addressed, including its anaerobic nature, promoter sequences, antibiotic selection marker, functional replication systems for shuttle vectors, and more. In this study, we have made significant advancements in establishing genetic tools for *F. nucleatum*. Firstly, we d identified multiple promoters with varying strengths via a Nanoluc report system, providing prototypes for the development of promoters with different levels of activity. Additionally, we identified a new antibiotic marker, *aad9*, for spectinomycin selection, expanding the options for antibiotic selection in *F. nucleatum*. Through exploration of native plasmids in *F. nucleatum*, we established three different plasmid backbones, demonstrating compatibility between pFN1 and pKH9. Given the widespread use of dual-plasmid-based genetic tools in other bacteria (50–52), we established a dual-plasmid system for *F. nucleatum*, incorporating the newly identified *aad9* selection marker and the widely used *catP* marker, along with the newly identified compatible backbones. This well-designed dual-plasmid system lays the foundation for the development of valuable tools for studying this clinically relevant bacterium, enabling investigations into its gene expression patterns, environmental responses, and potential applications in dental and periodontal research.

## Materials and Methods

### Bacterial strains and growth conditions

*E. coli* strains were grown aerobically overnight at 37°C with LB (10 g/L NaCl, 5 g/L tryptone, 10 g/L yeast extract) broth or LB agar plate. *F. nucleatum subspecies nucleatum* ATCC25586 and *F. nucleatum subsp. polymorphum* ATCC10953 were obtained from the American Type Culture Collection (ATCC). *F. nucleatum* was cultured at 37°C in 80:10:10 (N_2_: H_2_: CO_2_) atmosphere on Columbia (2%) agar plate supplied with 5% sheep blood, referred as CBA plates. Liquid cultures used brain heart infusion (BHI) medium. Growth media was supplemented with antibiotics at the following working concentrations; chloramphenicol (15 µg/ml), thiamphenicol (5 µg/ml), spectinomycin (100 µg/ml). All solutions and plates were pre-reduced overnight in an anaerobic chamber to remove entrapped oxygen. The strains used in this study were provide in Table S1, and the oligos used were listed in Table S2.

### Construction of pFN1 and related plasmids

Plasmids used in this study are listed in Table S1. Plasmids were constructed using gel-purified DNA fragments generated by commercial restriction enzyme digestion or PCR products generated with a high-fidelity DNA polymerase (Phanta Master Mix; Vazyme) from plasmid or genomic DNA templates using oligonucleotide primers listed in Table S2. Details of individual plasmid constructs are described below. Initial plasmid constructs were isolated after transformation into *E. coli* DH5α. Plasmids used to transform *F. nucleatum* were propagated in *E. coli* ER2796. Recombinant plasmid constructs were verified by sanger sequencing and analyzed with SnapGene.

### Detection of DNA methylation by PacBio sequencing

The methylated DNA motifs of *F. nucleatum* ATCC25586 and ATCC10953 were identified by single molecule real-time sequencing (SMRT) with the PacBio Platform, the generated data was deposited at both NCBI and the NEB rebase database and the details are descripted at the data availability section. The genomic DNAs of *F. nucleatum* ATCC25586 and ATCC10953 was isolated with gDNA isolation kit (Vazyme). A 10K SMRT Bell library was constructed using the SMRTbellTM Template Kit (version 1.0) reagent kit. The DNA samples, which passed the electrophoresis test, were sheared into fragments of the required size for library construction using Covaris g-TUBE. After DNA damage repair and end repair, hairpin adapters were ligated to both ends of the DNA fragments using DNA ligase, and the DNA fragments were purified using AMpure PB magnetic beads. Specific-sized fragments were selected using BluePippin fragment screening, and the SMRT Bell library was subjected to concentration selection using AMpure PB magnetic beads. Subsequently, DNA damage was repaired, and the SMRT Bell library was purified again using AMpure PB magnetic beads. The constructed library was quantified using Qubit, and the insert fragment size was detected using the Agilent 2100. Finally, sequencing was performed using the PacBio platform by Novagene (Beijing, China). The DNA methylation sites and motifs were analyzed with SMRT Link v5.0.1, which can predict 6-mA, 4-mC, 5-mC and unknown modified_base.

### Multiple amino acid sequence alignment and domain analysis of FNP_0020

The FNP_0020 protein sequences were retrieved from NCBI (https://www.ncbi.nlm.nih.gov/pro-tein) and used in multiple sequence alignment and domain analysis. In total, 46 genomic files of *Fusobacterium nucleatum* subspecies, along with 282 proteome files of *Fusobacterium* species were downloaded from the NCBI database (up to January 2024). Multiple amino acid sequence analysis was performed with Clustal X 2.1 using complete sequences from the selected prokaryotic species. Phylogenetic tree was constructed using “Draw tree” option in Clustal X using NeighbouJoining algorithm with default settings (Gap opening:10, Gap extension:0.2, bootstrap number:1000). The out-put PHYLIP file was analyzed using interactive Tree of Life (iTOL) software for scientific representation.

We queried Entrez (https://www.ncbi.nlm.nih.gov/cdd/) to access the CDD’s domain information in the CDD resource about FNP_0020 protein. The conserved domain summary pages give access to a wealth of data associated with each domain family, including hierarchical classifications, taxonomic information, sequence alignments, structural interaction data, domain architectures, functional site annotations, and literature.

### Identification and classification of *F. nucleatum* DNA methyltransferases

We combined the results of PacBio SMRT sequencing and the bioinformatic analysis from REBASE database. The REBASE database provided the predicted information about the restriction modification systems of both *F. nucleatum subsp. nucleatum* ATCC25586 and *F. nucleatum subsp. polymorphum* ATCC10953.

### Cloning and expression of MTases in *E. coli*

The MTase encoding genes of *F. nucleatum* ATCC25586 and ATCC10953 were firstly codon-optimized for suitable expression in *E. coli* and then synthesized as gBlocks by Sangon Biotech, China. The synthesized gBlocks were amplified by PCR and cloned into plasmid pSZU211 using Gibson Assembly. The primers used to amplify the gBlocks and the vector backbone were listed in the primer list (Table S2). Three genes encoding the MTases, including FN2074, FN1935 and FN1923 for *F. nucleatum* ATCC25586, FN1716 and FN1923 for *F. nucleatum* ATCC10953, were inserted into pSZU211 backbone arranged as co-transcription using Gibson Assembly. The pSZU211 inserted with MTases genes was confirmed by sanger sequencing and then conjugated into intermediatory *E. coli* hosts.

Tri-parental conjugation was employed to integrate the pSZU211 inserted with FN2074, FN1935 and FN1923 onto chromosome of *E. coli* ER2796 (Dam-), and the pSZU211 inserted with FN1716 and FN1923 onto chromosome of *E. coli* DH5α (Dam+). The procedures for tri-parental conjugation were developed based on the previously reported protocol (citation, pMID:). All the *E. coli* strains, including helper strain SZU195, donor strains SZU604 and SZU693, and recipient strains ER2796 and DH5α were cultured in LB broth supplied with appropriate antibiotics or 300 µM DAP at 37℃, 220 rpm until OD_600_ reached 0.5∼0.6. All the cultures were washed once with LB fresh medium by centrifugation at 4000 g, 5 min, and resuspended with equal volume of LB medium. After mixing 1 ml each of helper, donor, and recipient cultures, the bacterial resuspension was centrifuged at 4000 g, 5 min. The resulting bacterial pellets were resuspended with 30 µl of LB medium and spotted onto LB agar plates supplied with 300 µM DAP. The plates were then incubated for 18 hours at 37°C. After incubation, the mating lawns were resuspended in 900 µl of PBS and plated onto LB agar plates containing 20 µg/ml gentamycin for selection. Successful transconjugants were confirmed by PCR and sanger sequencing.

### Identification of MTases responsible for DNA methylation motifs

To idetermine the methyltransferase responsible for each identified DNA methylation motif, several intermediate E. coli strains expressing individual putative methyltransferases were constructed. For *F. nucleatum* ATCC25586, FN1923, FN1935, FN2071, and FN0416 were tested individually, whereas for *F. nucleatum* ATCC10953, FN1923 and FN1716 were tested. Meanwhile, the *E. coli* MTase YhdJ was included as a control for this assay. Cloning and integration of these genes into *E. coli* ER2796 were performed in the same way as described in the above section. Note that E. coli ER2796 was used as the host in this assay, as it is a MTase-free strain, to avoid interference from the intrinsic MTases of the *E. coli* host.

Methylation sensitive endonuclease BspHI, EcoRV, FokI and AcuI were employed to distinguish the methylated and unmethylated DNA motifs on the plasmids. Plasmids with the identified DNA methylation motif were selected for this assay, including pKH9-RM and pAsCas12f. The DNA methylation motifs on these plasmids were designed in a way that incorporates a mosaic of both DNA methylation recognition sequences and endonuclease recognition sequences. The enzymes and plasmids used for identification of the DNA motifs of each putative MTases were listed in Table S3. The plasmids were firstly transformed to the *E. coli* ER2796 or *E. coli* DH5α expressing individual or multiple MTase genes, and then isolated for subsequent restrict digestion. Note that plasmids isolated from native *E. coli* ER2796 or *E. coli* DH5α were included as unmethylated controls. Methylation-sensitive endonuclease digestion was performed according to the manufacturer’s protocol for each enzyme, followed by analysis of the digestion products using DNA gel electrophoresis. Note that the correct MTase for the DNA motif can protect the DNA from digestion, which can be observed in the electrophoresis results.

### Cloning and expression of FNP_0020 in *E. coli* ER2796

The FNP_0020 encoding genes of ATCC10953 were firstly codon-optimized for suitable expression in *E. coli* and then synthesized as gBlocks by Sangon Biotech, China. The synthesized gBlock was amplified by PCR and cloned into plasmid pSZU211 with LacI fragments using Gibson Assembly. The primers used to amplify the gBlocks and the vector backbone were listed in the primer list (Table S2). FNP_0020 and LacI fragments were inserted into pSZU211 backbone using Gibson Assembly. The pSZU211 inserted with FNP_0020 gene was confirmed by sanger sequencing and then conjugated into E. coli ER2796 via tri-parental conjugation as described above.

### Determination of FNP_0020 responsible for GATC motifs methylation

To determine the FNP_0020 responsible for GATC motifs methylation, *E. coli* strain expressing individual FNP_0020 were constructed and were tested. Meanwhile, the *E. coli* ER2796 (Dam-) was included as a control for this assay. Cloning and integration of FNP_0020 gene into E. coli ER2796 were performed in the same way as described in the above section. Note that E. coli ER2796 was used as the host in this assay, as it is a MTase-free strain, to avoid interference from the intrinsic MTases of the E. coli host. Specific Dam+ endonuclease DpnI were employed to distinguish the methylated and unmethylated DNA motifs on the plasmids. Plasmid pFN1-P4.5s-Nanoluc with the identified DNA methylation motif were selected for this assay. The plasmids were firstly transformed to the E. coli ER2796 expressing FNP_0020 gene, and then isolated for subsequent restrict digestion. Note that plasmids isolated from native E. coli ER2796 were included as unmethylated controls. Specific endonuclease DpnI digestion was performed according to the manufacturer’s protocol for each enzyme, followed by analysis of the digestion products using DNA gel electrophoresis. Note that the correct FNP_0020 for the DNA motif can lead to the DNA cleavage from digestion, which can be observed in the electrophoresis results.

### Methylation of plasmids via the engineered intermediary *E. coli*

Various shuttle and suicide plasmids were firstly constructed with commercial *E. coli* DH5α competent cells. The plasmids were then isolated and transformed into the *E. coli* ER2796 or *E. coli* DH5α strains carrying MTase encoding genes. The MTases were constitutively expressed in these *E. coli* hosts, enabling methylations of the DNAs inside the cells. The plasmids were isolated from the intermediary *E. coli* hosts prior to transformation into *F. nucleatum* ATCC25586 or ATCC10953.

### Preparation of *F. nucleatum* competent cells and electroporation

The procedures for preparation of *F. nucleatum* competent cells and electroporation were similar to a previously reported protocol (10) with minor modifications. Specifically, *F. nucleatum* ATCC25586 and ATCC10953 were streaked from a −80℃ frozen stock onto CBA plates followed by 2-3 days of incubation in an anaerobic chamber at 37°C to obtain single colonies. After that, about 10 of colonies were collected and inoculated into 12 mL of BHI and cultured overnight at 37°C in an anaerobic chamber. After 2-3 days, typically 2 days for *F. nucleatum* ATCC10953 and 3 days for *F. nucleatum* ATCC25586, transfer the entire 12 mL of the overnight culture into 40 mL of fresh BHI in a 50 mL centrifuge tube. Grow cells at 37°C in an anaerobic chamber for 3-4 days, typically 3 days for *F. nucleatum* ATCC10953 and 4 days for *F. nucleatum* ATCC25586, until the OD_600_ reaches 1.2. The culture was then centrifuged at 6000-7000

### NanoLuc luciferase assay

The NanoLuc luciferase assay was performed as previously described (53). *F. nucleatum* strains were grown to mid-exponential phase (OD_600_ = 0.5∼0.6). 100 µL of the culture was mixed with 10 µL with the NanoLuc Reaction Buffer of the Nano-Glo Luciferase Assay System (Promega). Luciferase activities were measured every 2 minutes with a Tecan Spark microtiter plate. Luciferase values were normalized by the OD_600_.

### Western blotting of FomA mutants from *F. nucleatum*

Bacterial proteins were extracted using whole protein extraction kit (Cat.KGP250) by sonication. Protein concentrations were quantified by BCA Protein Assay Kit (Cat.P0012). Samples were separated using 10% SDS-PAGE and transferred to PVDF membrane (Millipore). Membrane was blocked for 30 min in 5% milk in TBS-0.1% Tween and incubated with the primary antibody against FomA overnight at 4℃. After washing three times with TBST, the membrane was incubated with HRP-conjugated Goat Anti-Mouse second antibody (Cat.S0002) at room temperature for 1 h. Blots were visualized using Enhanced Chemiluminescence kit (Cat.KGP1128) by ChemiDoc ™ XRS+ Imaging System (Bio-Rad).

### Delivery of DNA via *E. coli - F. nucleatum* conjugation

The *F. nucleatum* ATCC10953 was streaked from a −80℃ frozen stock onto CBA plates, followed by incubation in an anaerobic chamber at 37°C for 2 days to obtain single colonies. Two to five colonies were collected and inoculated into 4 mL of BHI medium followed by incubation at a 37°C anaerobic chamber for about 36∼48 hours until the OD_600_ reaches 0.5∼0.6. The donor strain *E. coli* WM3064 with the conjugative plasmid was cultured in LB broth with 15 µg/ml chloramphenicol and 300 µM diaminopimelic acid (DAP) at 37°C, 220 rpm. The overnight culture of donor strain was sub-cultured by 1:100 dilution into fresh LB broth with antibiotic for about four hours before conjugation, when the culture reached an OD_600_ between 0.4∼0.5. After that, the donor culture and recipient culture were transferred into the same 1.5 ml microcentrifuge tube with various ratios, including donor:recipient as 1:1, 1:2, 1:4, 2:1, and 4:1. For example, for a 1:1 ratio, 100 μl of donor culture was mixed with 100 μl of recipient culture. For a 1:2 ratio, 100 μl of donor culture was mixed with 200 μl of recipient culture. Then, fresh BHI medium was added to the same tube to increase the final volume to 1 ml. The diluted donor and recipient mixture was centrifuged at 7000 *g* for two minutes, followed by washing with 1 ml of BHI. Two negative controls were established: the donor-only and recipient-only groups. In these controls, the mixture was replaced with pure donor bacteria or pure recipient bacteria, respectively. The bacterial pellet was then resuspended with 30 μl of BHI by gentle pipetting. The resuspended mixture of donor and recipient bacteria was spotted onto CBA plates with 300 µM diaminopimelic acid followed by incubation at 37°C under anaerobic condition for 24 hours. After the incubation period, the bacteria on the agar plate were harvested using an inoculation loop and transferred into 200 µl of PBS. The bacterial pellets were thoroughly resuspended before plating onto CBA plates supplemented with 5 µg/ml thiamphenicol for selection. The plates were cultured at 37°C under anaerobic condition for 48-72 hours.

### Fluorescent Microscopy

*F. nucleatum* strains, including the wild-type strain and SZU1387, SZU1388 and SZU1389 were were grown in BHI, with appropriate antibiotics, to mid-exponential phase (OD600∼0.6 to 0.7). One milliliter of the bacterial culture was spun down and washed twice with PBS (pH=7.4) by centrifugation at 8000 g, 2 min. The bacterial pellets were then resuspended with 40-100 µl of PBS. The bacterial resuspensions were then spotted onto 1% agarose slides for fixation. Microscopy was then performed using a Zeiss Imager.Z2 fluorescent microscope with 100× oil immersion objective. To visualize the green fluorescence, the Alexa Fluor 488 channel was used, with exposure time of 100 ms. The acquired images were processed via ImageJ (https://imagej.net/software/imagej/).

## AUTHOR CONTRIBUTIONS

Experimentation: LL, YZH, TTZ, YMH, RG, MYL. Study design and analysis: LL, TTZ, XL, HWZ. Writing – original draft: LL and XL. Writing – review & editing: LL, TTZ, YZH, XL, HWZ.

## DATA AVAILABILITY STATEMENT

Materials are available upon reasonable request with a material transfer agreement with SZU and SMU for bacterial strains or plasmids. The plasmids used in this study were shared via Wekwikgene (https://wekwikgene.wllsb.edu.cn/). Plasmid sequence and map can be accessed on a Benchling link at https://benchling.com/feedbackkk/f_/z4APHZVZ-f-nucleatum-expressing-plasmids/. The assembled genomes of *F. nucleatum* ATCC25586 (assession: CP144203) and ATCC10953 (accession: CP144201 for genome, CP144202 for plasmid) were uploaded at NCBI under BroProject accession number PRJNA1044226 and PRJNA1051802, respectively. The Pacbio sequencing data were deposited at the Rebase database and can be visited via the link (http://rebase.neb.com/rebase/private/pacbio_Liu30.html).

## ACKNOWLEDGMENTS

This work was supported by National Key Research and Development Program of China (2023YFD 10800100) to X.L; National Nature Science Foundation of China grant 82270012 (X.L), 81925026 (HWZ), 82341218 (HWZ); the Science and Technology Project of Shenzhen (JCYJ20220818095602006) and Shenzhen University 2035 Program for Excellent Research (86901-00000216) to X.L.

