## supplementary materials for "Equip *Fusobacterium nucleatum* genetic tool kits with compatible shuttle vectors and engineered intermediatory *E. coli* strains for enhanced transformation efficiency"

**Appendix for:**

**Enhancing Electro-Transformation and Conjugation Efficiency in *Fusobacterium nucleatum* via Complete Bypass of Restriction-Modification Systems**

Ling Liu^1#^, Yuzhang He^2#^, Tingting Zhang^2^, Yongmei Hu^2^, Mingyue Luo^2^, Hongwei Zhou^2^*, Xue Liu^1^*

^1^ Microbiome Medicine Center, Department of Laboratory Medicine, Zhujiang Hospital, Southern Medical University, Guangzhou, Guangdong, China.

^2^ Department of Pathogen Biology, Base for International Science and Technology Cooperation: Carson Cancer Stem Cell Vaccines R&D Center，International Cancer Center，Shenzhen University Medical School, Shenzhen, Guangdong, 518055, China.

^#^These authors contributed equally

*Correspondence to

Hongwei Zhou:,

Or Xue Liu:

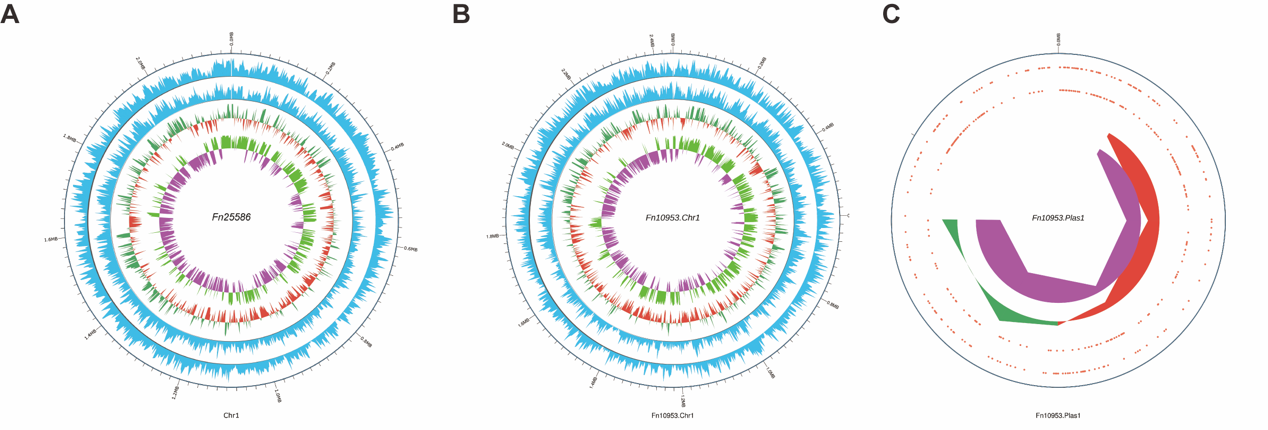

**Figure S1. Overview of DNA methylations of the genome within *F. nucleatum* ATCC25586 and ATCC10953.**

**A.** Genome modification within *F. nucleatum* ATCC25586. **B.** Genome modification within *F. nucleatum* ATCC10953. **C.** Genome modification within pFN3 plasmid from *F. nucleatum* ATCC10953.

The circles in the figure represent the following from the outermost to the innermost:

Genome position: Distribution of modification sites on the sense strand, calculated using a window size of 2000 bp and a step size of 2000 bp. Distribution of modification sites on the antisense strand, calculated using a window size of 2000 bp and a step size of 2000 bp.

GC content: Calculated using a window size of 2000 bp and a step size of 2000 bp. The inner red portion indicates regions with GC content lower than the average GC content of the entire genome, while the outer green portion indicates the opposite. Higher peaks indicate larger differences from the average GC content.

**Table S1. Strains and plasmids used in this study**

| **Strains/Plasmids** | **Description** | **Reference** |
| --- | --- | --- |
| ***F. nucleatum*** |  |  |
| ATCC 25586 | Wild type | (1) |
| ATCC 10953 | Wild type | (2) |
| SZU1207 | ATCC 25586, /pFN1-GFP, *catP^R^* | This study |
| SZU1378 | ATCC 25586, /pFN1-aad9, *spec^R^* | This study |
| SZU1379 | ATCC 25586, /pKH9-catP, *catP^R^* | This study |
| SZU1380 | ATCC 25586, /pFN1-aad9, pKH9-catP, *spec^R^*, *catP^R^* | This study |
| ***E. coli*** |  |  |
| ER2796 | λ-*fhuA2* ∆*(lacZ)r1 glnV44* *mcr-62* *trp-31* *dcm-6* *zed-501::Tn10 hisG1* *argG6 rpsL104 dam-16::Kan xyl-7* *mtlA2 metB1 (mcrB-hsd-mrr)114::IS10* | (3) |
| SZU1235 | ER2796, *glmS::Tn7* *FN1923* (Gm^R^) | This study |
| SZU1236 | ER2796, *glmS::Tn7* *FN1935* (Gm^R^) | This study |
| SZU1237 | ER2796, *glmS::Tn7* *FN2074* (Gm^R^) | This study |
| SZU1238 | ER2796, *glmS::Tn7* *FN0416* (Gm^R^) | This study |
| SZU1239 | ER2796, *glmS::Tn7* *FN1923 FN1935 FN2074* (Gm^R^) | This study |
| SZU1381 | ER2796, *glmS::Tn7* *C7Y58_05130* (Gm^R^) | This study |
| DH5α | *F− endA1 glnV44 thi-1 recA1 relA1 gyrA96 deoR nupG Φ80dlacZΔM15 Δ(lacZYA-argF)U169 hsdR17(rK− mK+) phoA λ–* | BBI LIFE SCIENCES |
| SZU693 | DH5α, *glmS::Tn7 FN1923 FNP1716* (Gm^R^) | This study |
| WM3064 | *thrB1004 pro thi rpsL hsdS lacZΔM15 RP4-1360 Δ(araBAD)567 ΔdapA1341::[erm pir]* | (4) |
| SZU195 | WM3064, expresses Tn7 transposase, *amp^R^* | (5) |
| SZU211 | WM3064, backbone for cloning Tn7L and Tn7R for transposons, *Gm^R^* | Lab |
| SZU603 | WM3064, /pFN1 -oriT, *Cm^R^* | This study |
| **Plasmids** |  |  |
| pCWU6 | *Cm^R^*; *Thia^R^* | (6) |
| pFN1-catP | *Cm^R^ /Thia^R^*, gBlock by BBI and Restriction-Methylation motifs silented from pCWU6 | This study |
| pFN1-Aad9 | *Spec^R^* | This study |
| pKH9-catP | *Cm^R^*; *Thia^R^* | This study |
| pKH9-RM | *Cm^R^*; *Thia^R^*, a plasmid was used for identifying the function of methyltransferases | This study |
| pFN1-AsCas12f | *Cm^R^*; *Thia^R^*, a plasmid was used for identifying the function of methyltransferases | This study |
| pFN1-P_4.5s_- NanoLuc | *Cm^R^*; *Thia^R^*, P_4.5s_ - NanoLuc | This study |
| pFN1- P_fomA_ - NanoLuc | *Cm^R^*; *Thia^R^*, P_fomA_- NanoLuc | This study |
| pFN1- P_fdx_ - NanoLuc | *Cm^R^*; *Thia^R^*, P_fdx_- NanoLuc | This study |
| pFN1-P_thl_ - NanoLuc | *Cm^R^*; *Thia^R^*, P _thl_ - NanoLuc | This study |
| pFN1-P_MT494_ - NanoLuc | *Cm^R^*; *Thia^R^*, P _MT494_- NanoLuc | This study |
| pFN1- P_pmhl_ - NanoLuc | *Cm^R^*; *Thia^R^*, P_pmhl_- NanoLuc | This study |
| pFN1- P_tetR_ - NanoLuc | *Cm^R^*; *Thia^R^*, P_tetR_- NanoLuc | This study |
| pFN1- P_xylR_- NanoLuc | *Cm^R^*; *Thia^R^*, P_xylR_- NanoLuc | This study |
| pFN1-catP-oriT | *Cm^R^*; *Thia^R^*, derivative of pFN1-catP with oriT | This study |
| pFN1-GFP | *Cm^R^*; *Thia^R^*, expressing GFP | This study |
| pKH9-mCherry | *Spec^R^*, expressing mCherry | This study |

Notes:

1. P_4.5s_ represents the promoter of 4.5s rRNA of *F. nucleatum*;

2. P_fomA_ is the promoter of fomA gene of *F. nucleatum*;

3. P_fdx_ is the promoter of ferredoxin gene of *Clostridium sporogenes*;

4. P_pmhl_ is the promoter of pGM-ACBQ plasmid used for *F. nucleatum*;

5. P_MT494_ is the promoter of ErmG from pMT494 plasmid used for *Bacteroides thetaiotaomicron*;

6. P_thl_ is the promoter of pGM-ACBQ plasmid used for *F. nucleatum*;

7. P_xylR_ is a xylose-inducible promoter and riboswitch combination system used for *F. nucleatum*;

8. P_tetR_ is the anhydrotetracycline-induce promoter of pRPF185 plasmid used for *Clostridium difficile*.

9. amp^R^: ampicillin resistance; Cm^R^: chloramphenicol resistance; Thia^R^: thiamphenicol resistance; Gm^R^: gentamycin resistance; specR: spectinomycin resistance.

**Table S2: Primers used in this study.**

| Primers | Sequences (5’— 3’) |
| --- | --- |
| 262_P18-mCherry backbone-F | CACCTGCCTCGcgccgcaaac |
| 263_P18-mCherry backbone-R | taattttcctcctCACCTGCC |
| 651_GN001800-F | GAGGCAGGTGaggaggaaaattaATGGAAAATTTGAAGCTGTTC |
| 721_p1800-backbone-R | gtgaaacctgctgatgtgctcTTACATTTCCTCTGTGTTGTAG |
| 722_A1812-F | gagcacatcagcaggtttcacacaggaaaactagtATGAATTATATTGGCTCAAAG |
| 723_A1812-R | ataaaactccTcCtccgcgcTTACTTTTTGATAAGACAATGC |
| 724_A1945-F | gcgcggaGgAggagttttatATGAGTTATAAGGAGAAGATCC |
| 725_A1945-R | gtttgcggcgCGAGGCAGGTGTTAATTTTTGATGAAATCATTGATGC |
| 1630_FNP1716-F | agcacatcagcaggtttcacacaggaaaactagtatgaaagaattttttaaaaatattg |
| 1631_FNP1716-R | gtttgcggcgCGAGGCAGGTGtcaaaacagactaaaatttttaacaac |
| 616_pCWU-ori-F | CTTCCGCTTCCTCGCTC |
| 617_pCWU-ori-R | CATGACCAAAATCCCTTAACGTG |
| 618_gBlock9-pKH9-Ori-R | GAGCGAGGAAGCGGAAGTGCTGGAGTGTGATATGTATGG |
| 619_gBlock9-pKH9-Ori-F | gactgccaggaaaattatttgtattc |
| 620_gBlock8-pCWU-catP-F | CACGTTAAGGGATTTTGGTCATGAGATTATCAAAAAGGATCTTCACC |
| 621_gBlock8-catP-R | attttcctggcagtcaattattt |
| 670_pKH9-catP-RMH-F | TGTctgatgccatgGGATGCTAGTAAAGTGTGAGAGCTTG |
| 671_pKH9-catP-RMH-R | CcatggcatcagACAGCTGTACTCTCTAGCATTAactaacc |
| 732_gBlock10-pFN1-F | GTTAGTAAAGTGTGAGAGCTTG |
| 733_gBlock10-pFN1-R | GAGCGAGGAAGCGGAAGTGATTTATTTTACTAGATATG |
| 734_gBlock8-catP-pFN1-R | CAAGCTCTCACACTTTAC |
| 952_pFN1-backbone-R | ctaagttccctctcaaattcaag |
| 953_pFN1-backbone-F | cttcaggtttgtctgtaactaaaaac |
| 1529_aad9-F | gaatttgagagggaacttagATGAATACATATGAACAGATTAATAAAG |
| 1530_aad9-R | agttacagacaaacctgaaggggattcatatggcTCATAATTTC |
| 950_pFN1-GFP-backbone-F | ggccaaccagataagtgaaatcGCGAGCGGTATCAGCTCACTC |
| 951_pFN1-GFP-backbone-R | gaattagcttcaaaagcgctcCGCAGCCGAACGACCGAGC |
| 187_lucFF-seqF | gagcgcttttgaagctaattc |
| 221_P18-reporter-SeqR | gatttcacttatctggttggcc |
| 193_FRT-attP-F | CGCCTTTGAGTGAGCTGATA |
| 973_4.5sRNA-F | TATCAGCTCACTCAAAGGCGgataatatatattgcccttaggggatatg |
| 974_4.5sRNA-R | atatatcctcctcactattttgaacataaagatacccctaggaag |
| 975_Nanoluc-F | caaaatagtgaggaggatatatatggtttttactctggaagattttg |
| 976_Nanoluc-R | ggaagataggcaattagtagaagc |
| 977_pFN1-Nanoluc-F | gcttctactaattgcctatcttccGTAATACGGTTATCCACAGAATCAG |
| 1358_pfomA-F2 | TATCAGCTCACTCAAAGGCGggagaaattttatattaaaatcaagaaaaaaagag |
| 1359_pfomA-R2 | caaaatcttccagagtaaaaaccatggttttttcccccttaaatttttg |
| 1521_Pfdx-F | TATCAGCTCACTCAAAGGCGgagaataggaacttcacgcg |
| 1522_Pfdx-R | tcttccagagtaaaaaccatacacttcctttccaactttatacg |
| 1523_pmhl-F | TATCAGCTCACTCAAAGGCGgtataaagttggaaaggaagtgtttg |
| 1524_pmhl-R | tcttccagagtaaaaaccatctaagttccctctcaaattcaag |
| 1525_pMT494-F | TATCAGCTCACTCAAAGGCGgtggtataatatctttgttcattag |
| 1526_pMT494-R | cttccagagtaaaaaccattataacctctccttaatttattgcatc |
| 1527_thl-F | TATCAGCTCACTCAAAGGCGGAgccatatgaatccctcctaat |
| 1528_thl-R | tcttccagagtaaaaaccattataacctctccttcacgcgtgaagttcctattc |
| 1143_Clo-TetR-R | aaaattttctcctttactgcaggag |
| 1282_pFN1-TetR-F | TATCAGCTCACTCAAAGGCGcaactactcccaacataaacatatattg |
| 1289_Nanoluc-F2 | gcagtaaaggagaaaattttatggtttttactctggaagattttg |
| 1290_Nanoluc-R2 | TCTGTGGATAACCGTATTACggaagataggcaattagtagaagc |
| 1360_Nanoluc-F2a | Atggtttttactctggaagattttg |
| 1290_Nanoluc-R2 | TCTGTGGATAACCGTATTACggaagataggcaattagtagaagc |
| 1361_Overlap-nanoluc-R | CTGATTCTGTGGATAACCGTATTAC |
| 1132_pFN1-catP-backbone-F | GTAATACGGTTATCCACAGAATCAG |
| 2160_xylO-F | TATCAGCTCACTCAAAGGCGGACATGATTACGAATTCGAGCTCT |
| 2161_xylO-R | tcttccagagtaaaaaccatCTTGTTGTTACCTCCTTAGCAGG |
| 1420_oriT-F | TATCAGCTCACTCAAAGGCGgtatactttccgctgcataaccc |
| 1421_oriT-R | TCTGTGGATAACCGTATTACctgttatgtttgggctcttcc |
| 2963_GFPoptFn-F | TATCAGCTCACTCAAAGGCGAAGCATAAAAAAAAGTTGGATAATATACTAAAATAG |
| 2963_GFPoptFn-F | GGGTTATGCAGCGGAAAGTAaaaaaagacttctcatgagagaagc |
| 2986_mCherry-F | gtggagtgtgaaccgagaccggtgagaatccaagcactagtaag |
| 2986_mCherry-F | ccatgccggcctgagaccaaaaaaaagacttctcatgagagaagcc |
| 2988_mcherry-backbone-F | ttggtctcaggccggcatgg |
| 2989_mcherry-backbone-R | ggtctcggttcacactccacaagc |

**Table S3 Restriction analysis of plasmid DNA with known endonucleases**

| MTase gene | Plasmid | Known endonuclease | Recognition sequence |
| --- | --- | --- | --- |
| FN1923 | pKH9-RM | FokI+ EcoRV-HF | TCATGA |
| FN1935 | pKH9-RM | BspHI+ EcoRV-HF | GGATG |
| FN2074 | pKH9-RM | BsiI | CCNNNNN/NNGG |
| FN0416  C7Y58_05130 | pKH9-RM | AcuI | CTGAAG |
| FNP0020 | pKH9-RM | - |  |
| FNP2222 | pKH9-RM | FokI + EcoRV-HF |  |
| FNP1716 | pFN1-AsCas12f | NsiI | ATGCATD |
